# Smartphone screening for neonatal jaundice via ambient-subtracted sclera chromaticity: *neoSCB* app pilot study

**DOI:** 10.1101/627034

**Authors:** Felix Outlaw, Miranda Nixon, Oluwatobiloba Odeyemi, Lindsay W. MacDonald, Judith Meek, Terence S. Leung

## Abstract

Jaundice is a major cause of mortality and morbidity in the newborn. Globally, early identification and home monitoring are significant challenges in reducing the incidence of jaundice-related neurological damage. Smartphone cameras are promising as colour-based screening tools as they are low-cost, objective and ubiquitous. We propose a novel smartphone method and app called neoSCB to screen for neonatal jaundice by imaging the sclera. It does not rely on colour calibration cards or accessories, which may facilitate its adoption at scale and in less economically developed regions. Our approach is to explicitly address three confounding factors in relating colour to jaundice: (1) skin pigmentation, (2) ambient light, and (3) camera spectral response. (1) The variation in skin pigmentation is avoided by imaging the sclera. (2) With the smartphone screen acting as an illuminating flash, a flash/ no-flash image pair is captured using the front-facing camera. The contribution of ambient light is subtracted. (3) This permits a device-and ambient-independent measure of sclera chromaticity following a one-time calibration. We introduce the concept of Scleral-Conjunctival Bilirubin (SCB), in analogy with Transcutaneous Bilirubin (TcB). The scleral chromaticity is mapped to an SCB value. A pilot study was conducted in the UCL Hospital Neonatal Care Unit. 51 neonates were imaged using the neoSCB app and had a blood test for total serum bilirubin (TSB). The better of two models for SCB based on ambient-subtracted sclera chromaticity achieved r = 0.75 (p<0.01) correlation with TSB. Ambient subtraction improved chromaticity estimates in laboratory tests and screening performance within our study sample. Using an SCB decision threshold of 190μmol/L, the sensitivity was 100% (specificity 61%) in identifying newborns with TSB>250μmol/L (area under receiver operating characteristic curve, AUROC, 0.86), and 92% (specificity 67%) in identifying newborns with TSB>205μmol/L (AUROC 0.85). These results are comparable to modern transcutaneous bilirubinometers.

## Introduction

Smartphones present unique opportunities for healthcare thanks to their processing and sensing capabilities coupled with their ubiquity and connectivity. Researchers are investigating smartphone-based tools to track and manage ongoing conditions such as diabetes [1] and bipolar disorder [2], as well as point-of-care, one-time screening or diagnostic applications such as assessing pupil reflex after traumatic brain injury [3]. In some cases, smartphones are being used to replicate the functionality of specialised medical devices such as the heart rate and blood oxygenation measurements performed by pulse oximeters [4,5]. Many concepts involve using an external sensor or wearable to collect data which is then processed and stored on a connected smartphone. Others endeavour to use only the in-built sensors of the phone. For example, the in-built camera can be used to assess wounds [6], the in-built microphone to monitor lung function [7], and the in-built accelerometer to detect falls [8].

In this work we explore the possibility of screening for jaundice in newborns using a smartphone camera. Jaundice refers to the yellow discolouration of skin and sclerae caused by a build-up of bilirubin, a naturally occurring breakdown product of the haemoglobin in red blood cells. Our technique relies on quantifying the colour of the sclera, as the degree of scleral yellowing is indicative of the systemic concentration of bilirubin. In fact, the overlaying conjunctiva more readily accumulates bilirubin than the sclera [9]. Here we use ‘sclera’ as a shorthand for ‘sclera and conjunctiva’. The project goal is to achieve a screening performance comparable to that of commercially available transcutaneous bilirubinometers using only the smartphone itself (without any additional hardware or add-ons). To this end we introduce our screening app, which we call ‘neoSCB’ for ‘neonatal Scleral-Conjunctival Bilirubinometer’.

Jaundice can be caused by an increase in the rate of haemolysis or a decrease in the rate of bilirubin excretion. A majority of newborns are affected by jaundice due to their immature livers and a high load of fetal haemoglobin breakdown. Other factors such as preterm birth and certain genetic disorders can significantly increase the risk of severe jaundice. When a neonate is severely jaundiced for an extended time, a neurotoxic form of bilirubin crosses the blood-brain barrier and can lead to permanent neurological dysfunction (kernicterus) or death [10].

Early identification is crucial: once diagnosed, jaundice can be effectively treated using phototherapy or, in extreme cases, exchange transfusion. In hospitals, newborns suspected of being at risk can be given a blood test. This is the gold standard diagnostic technique for determining the concentration of bilirubin in the blood. However, jaundice may only become problematic some weeks after birth, by which time the mother and newborn have typically been discharged. Screening for jaundice in the home environment presents several challenges, and current screening methods have inherent limitations. The most reliable screening tool is the transcutaneous bilirubinometer (TcB), a point-of-care device that takes a contact-based optical measurement of the skin to estimate bilirubin concentration. However, this device is expensive and not widely available, especially in less economically developed countries. The next most reliable screening approach is a visual examination by a trained healthcare professional, who can also look for behaviours associated with jaundice. Unfortunately, visual identification of jaundice is unreliable, with significant inter-observer variation [11,12]. The UK’s National Institute for Health and Care Excellence (NICE) explicitly recommends against using visual inspection as the sole means of screening [13].

In low-resource and remote areas, it may be that neither TcB screening nor expert visual assessment are options. In such cases, new parents may suffer anxiety and unnecessary, costly trips to the hospital. In the worst cases, severe jaundice goes unnoticed and leads to permanent disability or death.

The global burden of jaundice was estimated in 2010 to amount to 114,100 deaths and 75,400 cases of kernicterus each year [14]. The regions worst affected by jaundice-related death and disability are Sub-Saharan Africa and South Asia. These two regions account for three quarters of global mortality. Long-term impairments from kernicterus such as hearing loss and cerebral palsy are an order of magnitude more common in Sub-Saharan Africa than in developed countries [14]. Reasons for this include a lack of contact with healthcare professionals, a greater prevalence of certain genetic risk factors, such as G6PD deficiency, and the difficulty of spotting yellow discolouration in newborns with darker skin. The need for an accessible and reliable means of screening is particularly acute in these regions.

A smartphone camera-based screening method promises to be both objective and contactless. Importantly, smartphones are now ubiquitous even in resource-poor parts of the world. The impact of turning these devices into globally accessible home screening devices would be significant.

Research interest in smartphone-based jaundice screening is growing, and recent work is promising [15–22]. The most extensively validated system is the BiliCam app that estimates total serum bilirubin (TSB) based on the skin colour of the sternum [15]. Taylor et al. showed BiliCam performed at least as well as the latest models of TcB over a multi-ethnic sample of 530 newborns [18]. BiliCam uses a custom calibration card to standardise measurements, and machine learning on six images to estimate the TSB.

neoSCB differs in two main respects. Firstly, we image the sclera rather than the skin. Leung et al. compared sclera and skin regions using digital photography and found a stronger correlation between TSB values and sclera colour [23]. The sclera accumulates bilirubin while lacking the melanin and haemoglobin chromophores found in skin. It has been suggested that this permits a more sensitive measure of jaundice [17,23]. Secondly, we introduce a novel ambient subtraction method using the screen of the phone to illuminate the subject. This technique removes the need for a calibration card in the scene by explicitly subtracting the influence of ambient light on the measurement. This is important as the need for a calibration card may represent a significant barrier to adoption in a global context.

This paper introduces the neoSCB app and presents results from a pilot study testing the app in a neonatal population at University College London Hospital. One contribution of this paper is the ambient subtraction method using screen illumination, which serves as an equipment-free means of removing the variation due to ambient light. Another contribution is the demonstration that it is feasible to use this method in a clinical environment to image the newborn sclera. A third is the validation of the predictive utility of the Jaundice Eye Colour Index (JECI) proposed in a previous publication [24], and the comparison of this metric against another sclera-chromaticity-based TSB prediction model. We also introduce the concept of a Scleral-Conjunctival Bilirubin (SCB) measurement, analogous to the commonly used Transcutaneous Bilirubin (TcB) measurement.

This paper is structured as follows:

In the ‘Theoretical background’ section, the process of image formation, the motivation for the ambient subtraction method, and the concept of a device-independent colour space are outlined.
In ‘Methods’, the development and functioning of the neoSCB app is described, and an experiment in a controlled environment shows that ambient subtraction can improve the chromaticity estimate of a range of colours typical of a jaundiced eye. The clinical data collection procedure is explained. The processing of the image data to arrive at TSB estimates via sclera chromaticity estimates is outlined.
In ‘Results’, we compare a simple linear regression model based on the JECI metric to a multiple linear regression model based on chromaticity. We quantify the screening performance in our study population.
In ‘Discussion’, we summarise the challenges in attaining a reliable colour descriptor, and how our method seeks to address them. The performance of the neoSCB app is compared to standard screening tools. We discuss the advantages and drawbacks of the approach, and some limitations of the pilot study.

## Theoretical background

### Image formation

When an object is imaged by an RGB digital camera, the signal recorded at each pixel sensor depends on the product of three independent factors: the spectral power distribution of the light incident on the object, the object’s reflectance, and the spectral sensitivity of the camera. For a given point on the object, Equation 1 shows the recorded value, *I*_*k*_, for the corresponding pixel sensor in terms of the incident light spectrum, *E(λ)*, the object’s spectral reflectance profile, *S(λ)*, and the relative camera sensitivity, *Q*_*k*_*(λ)*.

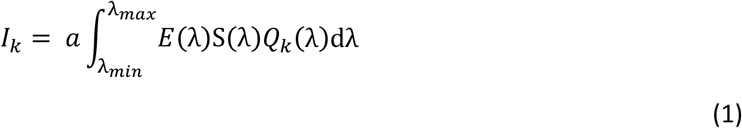

where *k* = *{R,G,B}* denotes the channel and *a* is a scale factor which accounts for geometry-dependent effects such as object shading, as well as scaling due to camera optics and sensor efficiency. The integral limits are determined by the sensitivity range of the camera.

To extract useful information about the object properties, one must isolate spectral reflectance *S(λ)* from the other factors that influence the recorded RGB values. This is problematic for several reasons. The light arriving at the camera sensor, sometimes called the colour signal *C(λ)*, is a product of two spectra, *C(λ)* = *E(λ)S(λ)*. It is impossible to isolate their individual contributions without contextual information; the same colour signal may result from two different objects given the right illuminations. Furthermore, the camera sensitivity functions *Q*_*k*_*(λ)* reduce a colour signal *C(λ)* to a single RGB triplet. This is a many-to-one operation: different colour signals may produce the same camera response. Finally, not all camera sensitivity profiles *Q*_*k*_*(λ)* are the same. There are variations between different makes and models, and even between different examples of the same device. This means that RGB triplets from different cameras cannot be directly compared, requiring conversion to a common colour space.

### Ambient subtraction

In most situations the ambient illumination spectrum is unknown. There is a large literature on methods to discount the effect of the ambient illumination on digital images [25–27]. Human vision exhibits colour constancy and so does this automatically. Image processing pipelines must first estimate the effect of the illumination and then scale the output RGBs values to remove it, such that a neutral object has equal RGB values. This is known as white balance. The aim of ambient subtraction is to circumvent the problem by providing an additional, known illumination, which we call the flash here. Two images are taken, one with flash and one without. By subtracting the output linear RGB values of the second image from the first image, it is possible to estimate the image as it would have been recorded under only the flash illumination. White balance can then be performed using *a priori* knowledge of the camera and flash without the need for any estimation from image information.

The ambient subtraction technique is summarised by Equation 2 for a pixel at image coordinate *x*. 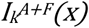 refers to the pixel value under the ambient and flash illuminations (from the first image captured), 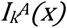 refers to the pixel value under only the ambient illumination (from the second image captured), and 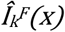 refers to the estimate for the pixel value under only the flash illumination.

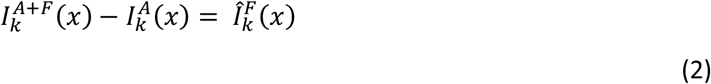

### Device-independent colour

A digital camera records RGB values according to its spectral sensitivity functions, which are device specific. These values are said to be in the internal or raw space of the camera. To be displayed or compared to other colour values, these raw triplets must be converted to a device independent colour space. A device independent colour space allows a description of a colour that is independent of the method of capture or display.

### CIE XYZ colour space

Based on the experiments of Wright and Guild in the 1920’s, the 1931 International Commission on Illumination (CIE) defined a standard observer that represents the average human response to colour stimuli within a 2-degree arc inside the fovea [28]. The XYZ space derived from this standard observer is a ‘reference’ colour space, meaning that it encompasses all the colours visible to the human eye. Most colour spaces are defined in relation to the CIE XYZ space, either directly or indirectly. It is often used as an intermediary space when transforming from a device-specific raw colour space to device-independent ‘output’ spaces, such as the commonly used sRGB space.

Developing a mapping between raw RGB values and XYZ values involves measuring or simulating a set of raw RGB values with known XYZ equivalents and then deriving a transformation matrix that minimises some colour error function. Typically, this is done using a specialised colour card with known XYZ values. Because illumination affects the colour of an object, the resulting transformation is only optimal for scenes under the same illumination used to develop the mapping. In general, no error free transformation from raw RGB to XYZ can exist because camera spectral sensitivity functions cannot be described as linear combinations of human cone sensitivities. In practice, however, 3×3 linear matrices suffice for most applications.

## Methods

### Ambient subtraction using screen illumination

The use of a screen as a source of illumination when using the front-facing camera was first implemented in smartphones by Apple Inc., with their so-called Retina Flash [29]. The neoSCB app uses the screen instead of the back-facing flash because the screen is not uncomfortably bright for the newborn. It provides an even and diffuse illumination when held near to the face.

neoSCB was developed for Android devices in the Android Studio IDE. It takes two raw format captures in succession using the front-facing camera. For the first, the screen is fully white and at maximum brightness. For the second, the screen is off. These will be referred to as the flash and no-flash images, respectively. The screen is white by default, turning off only briefly to take the second, no-flash image. This means the eye can adapt to the brightness, avoiding exposure to an over-stimulating flash. The image capture is triggered using the smartphone volume keys. Fig 1 shows the neoSCB app being used on a doll.

**Fig 1.**
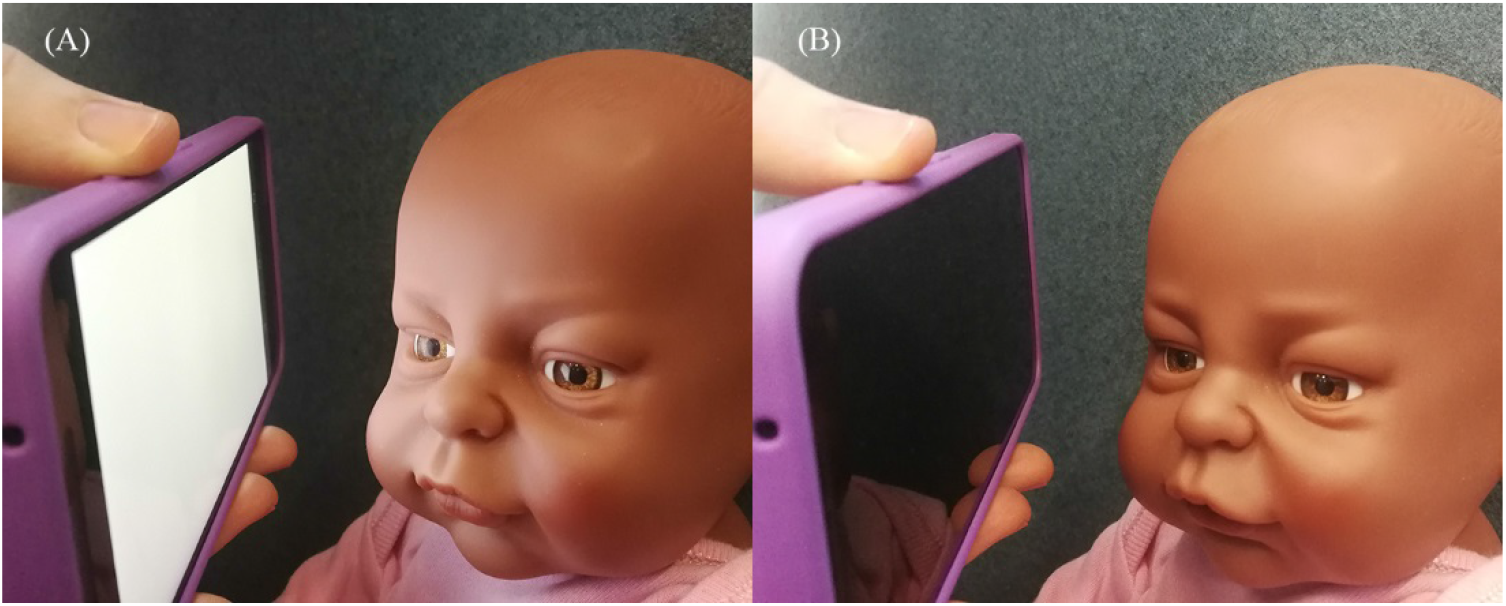
Demonstration of the use of the neoSCB app to capture flash/ no-flash sclera image pair. (A) Screen illuminates face for flash image capture. (B) Screen off for no-flash image capture.

There are several image processing techniques that rely on taking a flash and no-flash image pair of the same scene. These include estimating the ambient illumination, red eye removal, de-noising, and identifying foreground objects [30–32]. For our purposes, we are concerned with removing the influence of ambient light on the sclera colour recorded. By subtracting the colour values of the sclera under only the ambient illumination from the values obtained from the flash image, we can estimate the colour as it would appear under only the flash illumination. This ensures results are independent of the surrounding lighting environment.

### Data collection

51 neonates (35 male, 16 female) from the UCL Hospital Neonatal Care Unit outpatient clinic and Postnatal Ward were imaged using the neoSCB app over a 14-month period from January 2017. Images were captured at a convenient time during assessment and treatment. Typically, this was before the blood test while the baby was supine, but some images were taken while the baby was upright in the arms of a parent. All images were captured within 20 minutes of a blood draw for TSB determination. No additional blood tests that were not part of the routine care of the newborns were performed. None had received phototherapy within the preceding 24 hours. Written and verbal consent was obtained from all parents. Ethics approval was granted by the London – City Road and Hampstead NHS Research Ethics Committee. The study was conducted in accordance with the World Medical Association’s Declaration of Helsinki. The screen illumination was measured and confirmed to be safe for newborn eye (using appropriate aphakic hazard function) as per the International Commission on Non-Ionizing Radiation Protection Guidelines on Limits of Exposure to Incoherent Visible and Infrared Radiation [33].

Gestational age ranged from 35 weeks and six days to 41 weeks and one day. Postnatal age was distributed as follows: 17 subjects less than one week old; 24 subjects between one and three weeks old; 10 subjects greater than three weeks old.

The images were captured with a LG Nexus 5X smartphone held 10-20cm from the eye. The capture sequence took approximately one second. Multiple image pairs were captured when possible. Each image is saved in two formats: JPEG and DNG (a lossless raw image format by Adobe).

### Data analysis

Data analysis was performed in MATLAB R2018a (The Mathworks, Inc., USA). Fig 2 shows the processing pipeline applied to the images captured by the neoSCB app. It can be divided into two stages: first, the subtraction method is used to determine an xy chromaticity estimate for the sclera; next, a chromaticity-based prediction model is used to predict TSB. We call this TSB prediction from the sclera the Scleral-Conjunctival Bilirubin, or SCB, creating a concept analogous with the commonly used Transcutaneous Bilirubin, or TcB.

**Fig 2.**
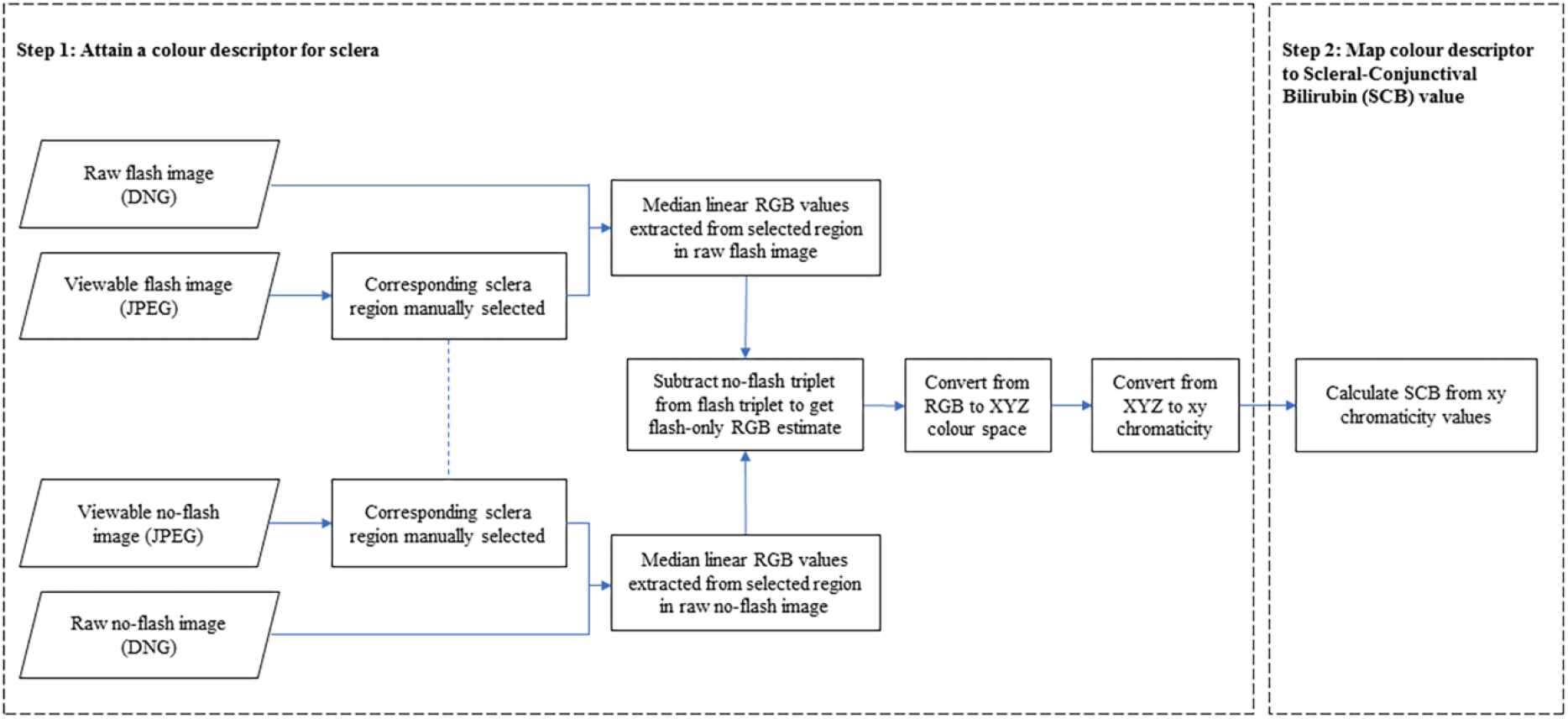
Processing pipeline to estimate scleral-conjunctival bilirubin from raw format flash/ no-flash image pairs. In Step 1, a color descriptor is estimated (ambient-subtracted sclera chromaticity). In Step 2, this is mapped to a scleral-conjunctival bilirubin (SCB) value.

### Chromaticity estimation via ambient subtraction

First, sclera regions of interest (ROIs) in the flash and no-flash images are manually selected. This is done on each image separately to allow for small movements between the two captures. Blood vessels and specular highlights are avoided, and same region of the eye is selected in each image. dcraw (v. 9.27) [34], an open source raw conversion software, is used to convert the DNG files into 16-bit, linear TIFF images. The images are minimally pre-processed: only black level subtraction and scaling to hardware saturation levels takes place. Using the ROIs and knowledge of the sensor Bayer pattern, the RGB values are extracted directly from the colour filter array output data. The median RGB values within the ROI are then calculated for flash and no-flash images. The median is used because it is robust when parts of the sclera ROI are not representative of the bulk colour. For example, part of a blood vessel or eyelash may be included by mistake.

The raw RGB values from the two images are then subtracted to provide an estimate of the raw RGB values as they would appear without ambient illumination. This subtraction is valid because the values are still in the linear colour space of the camera. If there are several image pairs available for one subject, each is used to generate a new ambient-subtracted RGB estimate. If there are several usable ROIs in a given image (for example, sclera regions either side of the iris), these are treated as independent measurements too. Sometimes, the signal from the flash is too low. This happens when the screen is not held close enough to illuminate the eye sufficiently, or when the ambient light signal is greater in the no-flash image. The latter can happen if there is movement between the flash and no-flash captures or the ambient light changes during image acquisition. Values that are less than one percent of the total bit depth after subtraction are discarded, as the signal from the flash is too low or the subtraction is invalid due to environmental changes. The final RGB estimate for a given subject is calculated as the median of the remaining post-subtraction RGB triplets generated. A linear 3×3 matrix transformation optimised for the screen illumination spectrum is then used to map these RGB values to device-independent XYZ values.

The phone-eye distance and absolute radiance of the phone screen are unknown, and therefore the subtracted XYZ values are only known up to a scale factor. Chromaticities (Equation 3), denoted with lowercase letters xyz, are the XYZ values normalised by the total (X+Y+Z). Chromaticities are invariant to anything that affects all channels equally, such as exposure time or phone-eye distance. By definition, the three chromaticities sum to one, and so only two chromaticity values are needed to fix the third: the chromaticity space is two-dimensional.

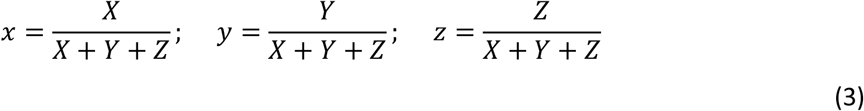

### Chromaticity-based models for estimating Scleral-Conjunctival Bilirubin (SCB)

We investigate two SCB models based on the estimated chromaticity values. The first, SCB_JECI_, is based on the Jaundice Eye Colour Index (JECI) proposed by Leung et al. [24]. JECI is defined in Equation 4 and was conceived as a proxy for the yellowness of the sclera. This SCB model is derived from a simple linear regression between JECI and TSB values. It assumes that the predictive utility of the chromaticity measurement is a function of how far along the yellow-blue axis the measurement is found. The JECI SCB model has one independent variable, so, necessarily, some chromaticity information is lost (chromaticity is two-dimensional). The second SCB model, SCB_xy_, uses both x and y chromaticity as independent variables. It is derived via a multiple linear regression of x and y chromaticity against TSB. The form of the SCB_JECI_ and SCB_xy_ models are shown in Equation 5 and Equation 6.

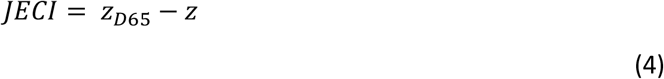

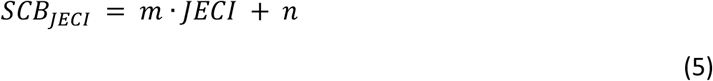

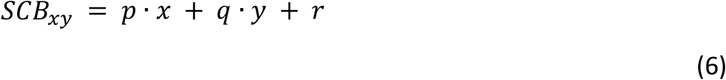

### Validation of ambient subtraction using screen as illumination

What follows is a demonstration that ambient subtraction can be used to improve chromaticity estimates under various ambient lightings. The test target was a printed grid of 11 yellow patches ranging from 0 to 0.1 on the JECI scale, where 0 JECI is white and positive values are yellow. A JECI value corresponds to a chromaticity value rather than a colour value, so the luminance Y was selected to be the largest that would fit within the printer gamut. The printed target was measured using a X-Rite Inc. ColorMunki Photo to determine a ground truth value for the test target chromaticities.

A TaoTronics TT-DL09 LED desk lamp was used to provide a controlled ambient light of two different correlated colour temperatures, 2700K (warm) and 6500K (cool). The smartphone was mounted at an angle of 45° and a separation 15cm from the test target centre for capture using the neoSCB app.

The pipeline shown in Step 1 of Fig 2 was used to determine chromaticity values from flash/ no-flash image pairs.

Fig 3 shows the results of the subtraction in the xy chromaticity space. For clarity, only data for every second patch is displayed. The human visual system chromaticity gamut is included for visual context. The ground truth chromaticities are plotted as filled circles in their respective colours (as measured using the ColorMunki). The empty circles represent the chromaticities measured under the no-flash condition, when the screen was off and only the lamp was illuminating the test target. The filled circles represent the chromaticities after the ambient subtraction. For both warm (Fig 3A) and cool (Fig 3B) ambient illuminations, the agreement is significantly improved by the subtraction method.

**Fig 3.**
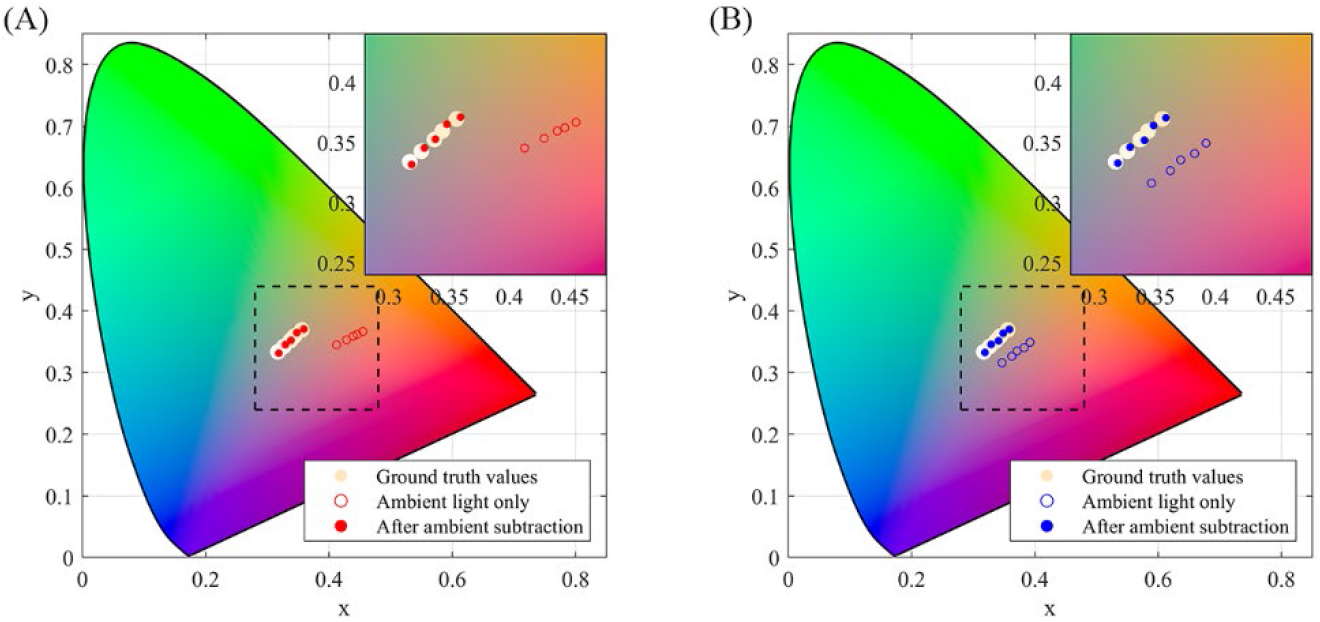
Visualising ambient subtraction in CIE 1931 chromaticity diagram for Jaundice Eye Colour Index (JECI) colours. JECI values are 0.01, 0.03, 0.05, 0.07, 0.09. After ambient subtraction, values come into alignment with ground truth values. (A) Under warm ambient illumination, correlated colour temperature is 2700K. (B) Under cool ambient illumination, correlated colour temperature is 6500K.

## Results

For two of the 51 subjects, no TSB was recorded. For a further 12, a flash/ no-flash image pair with the sclera visible was not successfully captured, or the post-subtraction values were less than the minimum threshold. SCB values were calculated for the remaining 37 subjects.

The correlation between TSB and SCB for the two SCB models are summarised in Table 1, alongside their root-mean-square residuals. A comparison of the models with and without the use of ambient subtraction shows that subtraction decreases RMS residuals and greatly increases correlation.

**Table 1.** Comparison of SCB models in terms of correlation with TSB and RMS residuals with/without ambient subtraction.

| SCB Model | Without Ambient Subtraction |  | With Ambient Subtraction |  |
| --- | --- | --- | --- | --- |
| | Pearson's r | RMS Residual ($\mu\text{mol/L}$) | Pearson's r | RMS Residual ($\mu\text{mol/L}$) |
| **SCB<sub>JECI</sub>** | *0.38 ($p > 0.01$)* | *70* | *0.70 ($p < 0.01$)* | *54* |
| **SCB<sub>xy</sub>** | *0.56 ($p < 0.01$)* | *63* | *0.75 ($p < 0.01$)* | *51* |

The best correlation is obtained using SCB_xy_, the multiple linear regression model for SCB, with ambient subtraction. Fig 4 shows the correlation between SCB_xy_ and TSB with and without ambient subtraction. Results under the white screen illumination show a correlation of 0.56 (p < 0.01). When the subtraction method is applied to suppress the effect of ambient light, the correlation increases to 0.75 (p < 0.01). The root-mean-square residual is 51 μmol/L.

**Fig 4.**
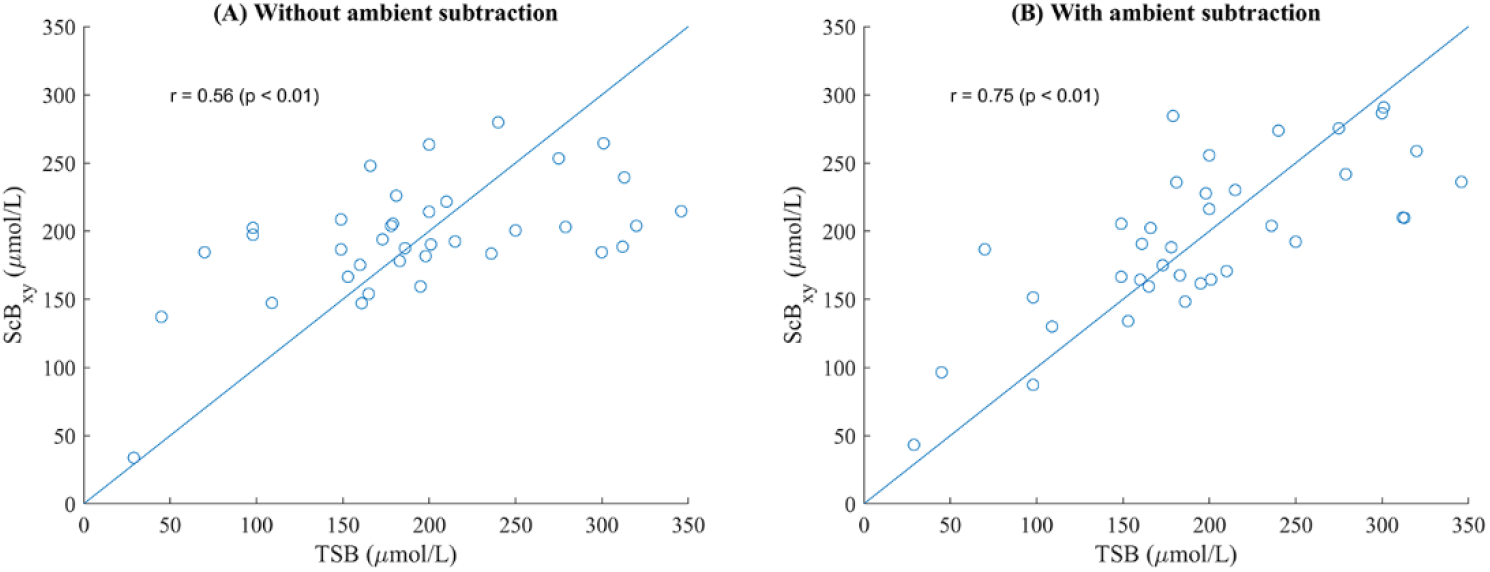
Scatter plot of scleral-conjunctival bilirubin (SCB) against TSB with and without ambient subtraction, 37 subjects. The use of ambient subtraction greatly improves the correlation. (A) Correlation without ambient subtraction, r=0.56. (B) Correlation with ambient subtraction, r=0.75.

Fig 5 displays the Bland-Altman plot for the ambient-subtracted SCB_xy_ method. The Bland-Altman plot shows that SCB_xy_ is systematically lower than TSB at high values, and vice versa. However, the important quantities are the sensitivity and specificity achieved as this technique is intended as a screening tool. Fig 6 shows Receiver Operating Characteristic curves for the ambient-subtracted SCB_xy_ method at two clinically relevant screening thresholds: 250μmol/L (Fig 6A) and 205μmol/L (Fig 6B). 250μmol/L is the threshold TcB measurement for babies greater than 35 weeks gestational age and older than 24 hours above which a blood test is recommended, and 205μmol/L is the treatment threshold for term babies at 24 hours, according to NICE guidelines [13].

**Fig 5.**
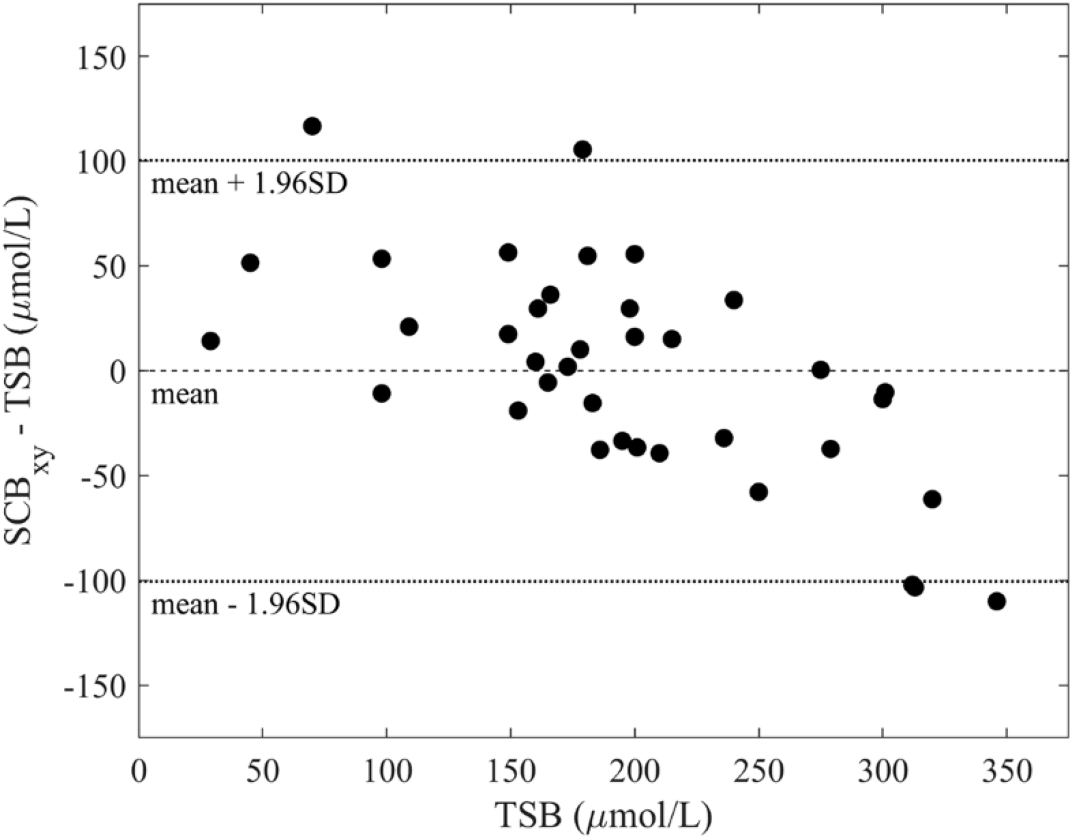
Bland-Altman plot for TSB and SCB_xy_ derived from ambient-subtracted scleral chromaticity.

**Fig 6.**
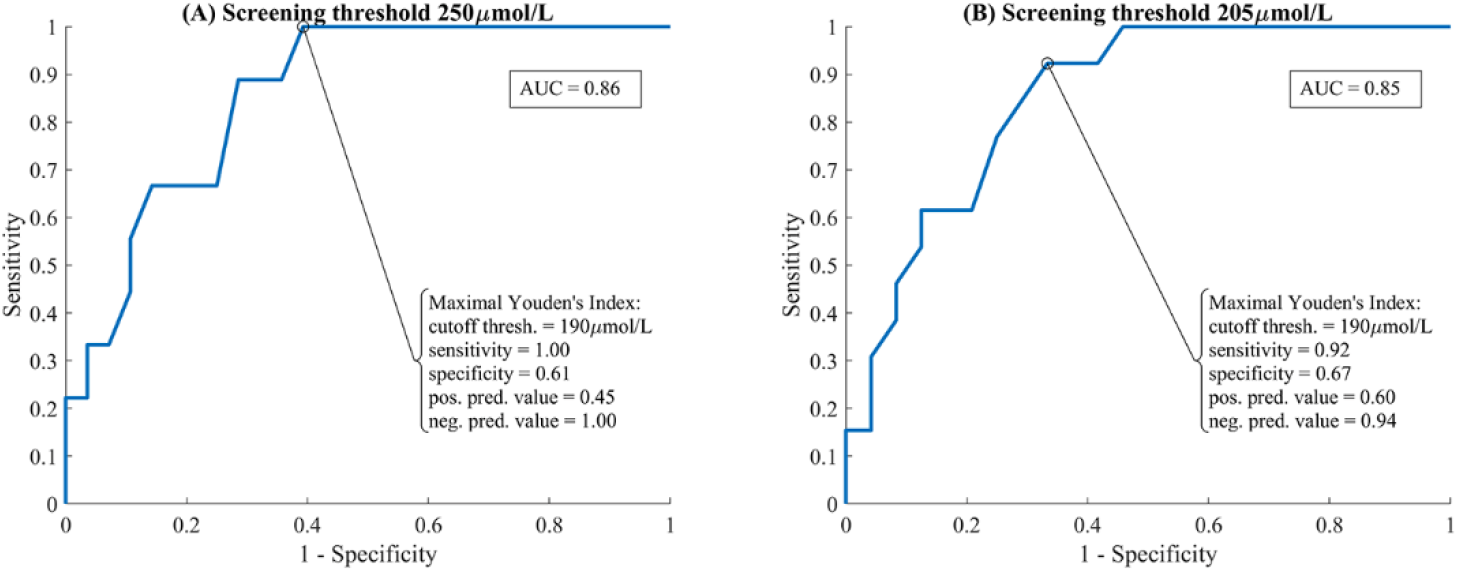
Receiver Operating Characteristic plots screening TSB at two thresholds using SCB_xy_ as the decision metric. (A) Screening threshold of 250μmol/L. (B) Screening threshold of 205μmol/L.

For a screening threshold of 250μmol/L, the area under curve (AUC) – which quantifies how effective the tool is at separating cases over and under the screening threshold – is 0.86 (a perfect tool would have AUC of 1, a random decision would have AUC of 0.5). The maximal Youden’s Index (true positive rate – false positive rate) is achieved for a cut off threshold of 190μmol/L. In this case, the screening method (with 95% confidence interval in brackets) has sensitivity 100% (88% - 100%), specificity 61% (43% - 76%), positive predictive value 45% (29% - 62%), and negative predictive value 100% (88% - 100%).

For a screening threshold of 205μmol/L, the AUC is 0.85. The maximal Youden’s Index is achieved for a cut off threshold of 190μmol/L. In this case, the screening method (with 95% confidence interval in brackets) has sensitivity 92% (78% - 99%), specificity 67% (49% - 81%), positive predictive value 60% (43% - 75%), and negative predictive value 94% (80% - 100%). To successfully classify every baby with TSB greater than 205μmol/L in our study sample, the SCB_xy_ cut off threshold is lowered to 170μmol/L. In this case, the screening method has sensitivity 100% (88% - 100%) and specificity 54% (37% - 70%).

## Discussion

neoSCB can be divided into two steps: first, attain a reliable colour descriptor for the tissue, and, second, map this to a SCB value. A screening decision can then be made based on the SCB value. In this work we have used the ambient-subtracted scleral chromaticity as our tissue colour descriptor, and a linear model to calculate SCB from it. This approach can be generalised: chromaticity could be mapped to SCB using more sophisticated models (the relatively small dataset prohibited this in our case), or a colour descriptor other than chromaticity could be used. Indeed, the linear model used to calculate SCB in this work systematically underestimates high TSB values, which could imply that a higher order model is required. Regardless of the SCB model used in step two, the performance will depend on the accuracy of the colour descriptor output by step one. In what follows we discuss some of the challenges in attaining an accurate colour descriptor and how they were addressed in this work.

### Attaining a reliable colour descriptor

There are several challenges in attaining a reliable and representative tissue colour descriptor by digital imaging: (1) Ambient illumination affects recorded colour (2) Devices may respond in different ways to the same light signal, both at the hardware and software level (3) The selected ROI should be ‘representative’: a specular region or a blemish or bleed is not a good area from which to measure the degree of yellowing due to hyperbilirubinemia.

### Discounting ambient illumination

To avoid the effect of ambient light, we have proposed a subtraction method. Our results indicate that the correlation between our chromaticity-based SCB models and TSB can be improved by subtracting the ambient component of the signal. There are several necessary requirements if ambient subtraction is to be effective. Crucially, the camera must have a linear response to light. Furthermore, no region of either image can be saturated, and both images must be captured with the same exposure time (or have values scaled to correct for this). The ambient light falling on the ROI cannot change in intensity or chromaticity between the flash and no-flash captures, otherwise the post-subtraction values will be spurious. Finally, ambient subtraction requires that the same ROI is visible in both images.

### Device independence

To ensure the technique is device independent, we capture in raw format to avoid the software-specific post-processing algorithms found in all smartphones. However, smartphone cameras can vary at the hardware level too because sensors have different spectral sensitivity profiles [35]. To mitigate this, we use a transform to a device-independent colour space, the CIE 1931 XYZ space.

### Avoiding specular reflection and ‘unrepresentative’ regions

By imaging the sclera, neoSCB avoids the variability introduced by different skin tones. However, despite the lack of melanin, the sclera colour signal may be influenced by factors other than bilirubin concentration. Non-representative and specular regions can sometimes be avoided by careful image segmentation. However, manual and automatic ROI selection system will both sometimes fail. To mitigate this source of error, multiple image pairs and regions can be analysed (with the assumption that only a minority of regions will suffer from these problems). Pixel values can be summarised with summary statistics robust to outliers such as the median.

### Comparison to transcutaneous bilirubinometers

With a sample size of 630, Romangoli et al. evaluated the BiliChek (Philips Healthcare) and Jaundice Meter JM-103 (Draeger Medical Inc.) transcutaneous bilirubinometers for a screening threshold of 205μmol/L [36]. The sensitivity and specificity achieved were 0.99 and 0.30, respectively, for BiliChek, and 1.00 and 0.42, respectively, for JM-103. neoSCB results are comparable for this screening threshold, with a sensitivity of 1.00 and a specificity of 0.54. However, neoSCB AUC was smaller at 0.85 compared to 0.89 for BiliCheck and 0.94 for JM-103.

In areas with poorer healthcare resources TcBs are not always available, even in hospital settings. Smartphone-based methods offer an accessible and cheap alternative that can provide an objective indicator of jaundice severity. Using smartphones could reduce the amount of equipment that visiting healthcare professionals need to transport, allow for home monitoring by parents, and increase the frequency of measurements in underequipped hospitals. In some hospitals blood tests are only available infrequently (e.g. once per day) and TcBs are not used. In these situations, neoSCB could be used to monitor response to treatment. It is also worth noting that TcBs are not used for babies undergoing phototherapy treatment, as the skin exposed is no longer thought to be representative of systemic bilirubin levels [37]. As the eyes are covered during phototherapy, this hypothesised bleaching effect may not affect the sclera colour. Another reason to prefer sclera colour measurement is the fact that it cannot be influenced by skin pigment. There is a body of evidence that TcBs overestimate jaundice levels in dark-skinned infants [38–41]. This may lead to unnecessary treatment which can block access to phototherapy beds and cause dehydration.

### Limitations and future work

One of the aims of the ambient subtraction approach is to minimise the need for equipment such as colour cards for calibration to lower the barrier to adoption. However, variation between the screen spectral power distributions and camera spectral sensitivities of different smartphone models means that some characterisation of the smartphone is needed to get a device-independent colour measurement. This would involve a one-time colour card characterisation performed by the user before use of the smartphone.

neoSCB relies on raw capture. Even if there is a trend towards allowing access to the raw values output by the front-facing camera, only a few phones currently meet this requirement.

The post-subtraction signal is determined by the proportion of the signal which comes from the flash. In this work, we discarded post-subtraction values less than one percent of the total bit depth during offline analysis. In a future version, an automatic system will warn users if the screen illumination is insufficient, requesting a less bright environment or the screen held closer to the face. In some cases, movement between flash and no-flash captures rendered the pair of images unusable. The capture sequence lasts less than one second, so this is not typically an issue.

Currently, segmentation of the sclera and thresholding of the post-subtraction pixel value results are both done after capture during an after-the-fact analysis. To provide a TSB prediction for the user at the time of capture would require the automation of these two steps. Sclera segmentation algorithms have been developed by researchers interested in biometric identification using sclera vasculature [42,43]. Mariakakis et al. developed their own segmentation using accessories as fiducials [17].

Although here we have processed images after the time of capture on a desktop computer, the pipeline is computationally simple enough to be performed in real time on the smartphone. This is advantageous because it removes the need for internet connection for cloud-based processing, which is often unavailable or too expensive for users in less economically developed regions.

Imaging the eye of the infant can be challenging due to unpredictable movements and the amount of time spent asleep. This study has shown that it is possible to collect this data using a smartphone in a time-constrained clinical environment. It was observed that infants typically open their eyes when feeding, and this can provide a suitable opportunity for measurement. In a home setting there is less time pressure than in a clinical assessment and so the measurement could be done at any convenient point.

Although sclera colour has less variation than skin colour, there can be confounding influences. Neonates can have a blue tint in the sclera because it is thinner than the adult sclera. Blood vessels and bleeds must be identified so they do not lead to spurious TSB estimates.

### Advantages of ambient subtraction for point-of-care colorimetry

The principle advantage of using the ambient subtraction approach is to remove the need of a calibration colour card present in the frame of every image. Especially in realistic settings in the home, in the clinic, and in the field, it can be difficult to position a colour card in the shot and almost impossible to keep it in pristine condition for an extended period of use. By applying an ambient subtraction, we can know *a priori* the effective illumination of the post-subtraction scene. This means the transformation to XYZ can be determined by a one-time calibration in controlled conditions and optimised to provide more accurate chromaticity information.

## Conclusion

In this work, we show that by explicitly addressing the confounding factors in colour measurement of jaundice (namely ambient light, camera characteristics, skin tone), a linear model based on sclera chromaticity can predict total serum bilirubin accurately enough to be useful as a screening tool. By employing a novel ambient subtraction technique using the screen as illumination, this can be done without the need for add-ons or colour calibration cards in the shot. Our app, neoSCB, is tested in a clinical environment, demonstrating that it is feasible to capture the necessary flash/ no-flash image pair of the newborn sclera using the smartphone’s front-facing camera. We have shown that the correlation between a simple colour metric (JECI) and the measured TSB can be much improved by ambient subtraction with this flash/ no-flash image pair.

We have proposed the concept of Scleral-Conjunctival Bilirubin (SCB), analogous to Transcutaneous Bilirubin (TcB). It is not yet clear whether the scleral-conjunctival or transcutaneous bilirubin concentration is a better proxy for serum bilirubin concentrations, or indeed whether serum or extravascular tissue concentrations are better predictors of neurological damage from bilirubin. We hope that by introducing the concept of SCB measurement further research into measuring jaundice via the sclera can be stimulated.

We have investigated the screening utility of two xy chromaticity-based SCB models. One based on the Jaundice Eye Colour Index (JECI) proposed by Leung et al. achieved a correlation of 0.70 (p<0.01) with TSB. Another based on a linear model of x and y chromaticity values achieved a correlation of 0.75 (p<0.01). Using the latter approach, we were able to achieve a sensitivity of 100% for a specificity of 61% when screening infants with TSB above 250μmol/L and a sensitivity of 100% for a specificity of 54% when screening infants with TSB above 205μmol/L, a performance comparable to that of transcutaneous bilirubinometers.

## Acknowledgements

The authors would like to thank all the study participants and their parents.

## Supporting information

**S1 Data. Clinical data for all 51 subjects**.

